# Advancing bee conservation in the US: gaps and opportunities in data collection and reporting

**DOI:** 10.1101/2023.12.05.569280

**Authors:** Josée S. Rousseau, S. Hollis Woodard, Sarina Jepsen, Brianne Du Clos, Alison Johnston, Bryan N. Danforth, Amanda D. Rodewald

## Abstract

Bee conservation in the U.S. is currently hindered by challenges associated with assessing the status and trends of a diverse group of >3000 species, many of which are rare, endemic to small areas, and/or exhibit high inter-annual variation in population size. Fundamental information about the distribution of most species across space and time, thus, is lacking yet urgently needed to assess population status, guide conservation plans, and prioritize actions among species and geographies. Using wild bee data from two public data repositories representing the contiguous U.S., we evaluated the availability and sufficiency of data for use in species assessments of wild bees. We also examined the number of bee species recorded in each U.S. state and the proportion of species with recent records (2012-2021). Although efforts to monitor bees continue to grow, there remains a massive paucity of data. Exceedingly few records (0.04%) reported both sampling protocol and effort, greatly limiting the usefulness of the data. Few species or locations have adequate publicly available data to support analyses of population status or trends, and fewer than half of species have sufficient data to delineate geographic range. Despite an exponential increase in data submissions since the 2000s, only 47% of species were reported within the last decade, which may be driven by how data are collected, reported, and shared, or may reflect troubling patterns of local or large-scale declines and extirpations. Based on our analysis, we provide recommendations to improve the quality and quantity of data that can be used to detect, understand, and respond to changes in wild bee populations.

## INTRODUCTION

Global evidence of wild bee declines has accumulated steadily over the last three decades. Although the conservation status for most of the world’s roughly 20,000 wild bee species has still not been assessed (Winfree, 2010; deMaynadier et al., 2023), the proportion of threatened species ranges from 12.5-45% of regional faunas among those groups that have been considered, such as the bumble bees (Cameron and Sadd, 2020; Bumble Bee Specialist Group, 2023) and a small proportion of other bee groups (NatureServe, 2023). Notably, certain bee groups, including pollen specialists with limited host plant associations, and species with larger body sizes and smaller phenological breadth, appear particularly susceptible to decline (Biesmeijer et al., 2006; Bartomeus et al., 2013; Hofmann et al., 2019; Bogusch et al., 2020). The putative drivers of wild bee declines are overwhelmingly anthropogenic and include widespread loss, degradation, and fragmentation of suitable habitat; non-target effects of broadly-deployed pesticides, such as neonicotinoids; and climate change, which has both direct influences, such as exceeding thermal limits, and indirect effects, such as reducing floral resource availability (reviewed in Potts et al., 2010; Goulson et al., 2015). Bee conservation is now a formal priority on the part of both national (e.g., Pollinator Health Task Force, 2015) and international efforts (e.g., Promote Pollinators, 2023). This is in part because bee-mediated pollination services are paramount to human food security and ecosystem stability (IPBES, 2016) and also due to the widespread recognition of the intrinsic value of biodiversity.

In the United States, there are over 3000 wild bee species, with a number of endemic and highly specialized species, and particularly high diversity found in arid lands of the Southwest (Meiners et al., 2019; Orr et al., 2021; Chesshire et al., 2023). Conservation of threatened bee species is carried out through both regulatory and non-regulatory governmental policies, as well as through a patchwork of voluntary, non-governmental efforts. At the Federal level, nine bee species, all within the genera *Hylaeus* and *Bombus*, are currently protected under the Endangered Species Act (U.S. Fish and Wildlife Service, 2016, 2017, 2021), with at least another five species being considered for listing (U.S. Fish and Wildlife Service, 2023). At the state level, species can be protected under state endangered species acts in cases when these laws apply to insects, such as in California, where four species of bumble bee are candidates for listing under the California Endangered Species Act (Sanders, 2022). However, the level of protection afforded by state endangered species acts varies greatly by state. Species can also be designated as conservation priorities as Species of Greatest Conservation Need (SGCN) through State Wildlife Action Plans (SWAPs) (Mawdsley and Humpert, 2016; deMaynadier et al., 2023), or regionally as sensitive species on US Forest Service and Bureau of Land Management managed lands. State Natural Heritage Programs, operating as part of the NatureServe network, can assign species ranks according to threat level, and these assessments inform state and regional lists of at-risk species. Federal to local-level conservation incentives can also stem from species assessments from the International Union for the Conservation of Nature, although at present only bumble bee assessments have been completed (Bumble Bee Specialist Group, 2023). Conservation action for rare and threatened pollinator species takes many forms, including a significant annual investment in pollinator habitat management and restoration, based on the premise that local pollinators are habitat-deficient and possibly declining. For example, a key goal of the U.S. Pollinator Health Task Force (2015) was to create or enhance >7 million acres of pollinator habitat by 2020. However, without a solid understanding of which species are declining, what those species need, and how populations respond to restoration, it is unclear whether conservation investments are actually improving outcomes for declining bee species. In the end, the ability to assess population status and evaluate the effectiveness of management interventions relies upon having sufficient data across species, locations, and time.

Two of the greatest limitations to understanding the full extent of wild bee declines in the U.S. are (i) widespread gaps in availability of (or access to) bee data (Orr et al., 2021; Chesshire et al., 2023), and (ii) lack of implementation of standardized collection protocols and practices, outside of their use for particular projects (Montgomery et al., 2020, 2021; Woodard et al., 2020). The lack of available bee data hinders our ability to adequately assess bee status and trends because there is a paucity of data for many species, regions, and time periods. Correspondingly, the extinction risk of most U.S. wild bees is unknown, with only ∼600 species, or <1/5 of the fauna, having been assessed according to criteria of the International Union for the Conservation of Nature (IUCN) or NatureServe, two of the most commonly used frameworks for species conservation status assessments. Among these 600 species, the ∼50 species in the genus *Bombus* have been assessed most thoroughly (Bumble Bee Specialist Group, 2023). Moreover, when wild bee species are assessed, they are likely to be determined to be “Data Deficient” or “Unrankable” because only limited data are available, and only species with very small ranges and known threats within those ranges are likely to be considered imperiled. The wild bee data that are available were overwhelmingly collected using unstandardized data collection protocols, and thus they are largely not interoperable or are difficult to analyze together in meaningful ways (Potts et al., 2010; Montgomery et al., 2020; Woodard et al., 2020). This precludes performing some analyses that are critical for conservation decision-making, such as estimating species ranges with species distribution models (SDMs) or calculating extinction risk through population viability analyses (PVAs). For example, occupancy models, which can be powerful for detecting trends while accounting for some of these issues, have only recently begun to be developed for bee species (Graves et al., 2020; Otto et al., 2021, 2023; Boone et al., 2023b, 2023a), and the data needed to calculate these models are not often collected in routine field surveys. Recent research from other groups of insects (especially butterflies) highlights declines in broadly distributed, formerly abundant species (e.g. Wepprich et al., 2019; Forister et al., 2021, 2023; Van Deynze et al., 2022). If similar declines are occurring in the wild bee fauna, as available assessments suggest, they are potentially going unnoticed.

Ideally, wild bee data collection, especially when it is carried out with the goal of supporting species conservation, would be performed to best accommodate the needs of the conservation entities who use these data to assess statuses and trends (Nichols and Williams, 2006; Carroll et al., 2023). These entities have data needs that are largely overlapping, despite some differences in the analyses they employ (Table 1). Importantly, assessments can be carried out even when only minimal, unstandardized data are available; however, when high-quality data sets are available for assessments, this can lead to much more meaningful and informative status assessments, benefitting bee conservation. Evidence of this can be found in recent work on two bumble bee species, *Bombus occidentalis* (Federal ESA listing status: petition is under a 12-month finding) and *Bombus affinis* (Federal ESA listing status: endangered as of 2017). *B. occidentalis*, which once had a broad distribution across the western U.S. (Milliron, 1971), now occupies only isolated pockets of its former range. Occupancy-based analyses for this species have leveraged both historical sampling efforts inferred from presence data and newly collected, standardized data. These analyses helped to both document the extent of decline (Graves et al., 2020) and identify the primary causes, particularly the role of neonicotinoid insecticides (Janousek et al., 2023). In the case of *B. affinis*, fully standardized surveys, where effort is known and not inferred, have been conducted in recent years to support occupancy modeling (Boone et al., 2023b, 2023a; Otto et al., 2023). This work has helped to optimize detection probabilities and ultimately improved monitoring program design, which is essential for efficient and effective monitoring. Both of these examples, in particular *B. affinis*, clearly demonstrate how data collection methods that are fully reproducible, and account for and report sampling effort and methods of data collection, empower wild bee conservation efforts.

**Table 1.**
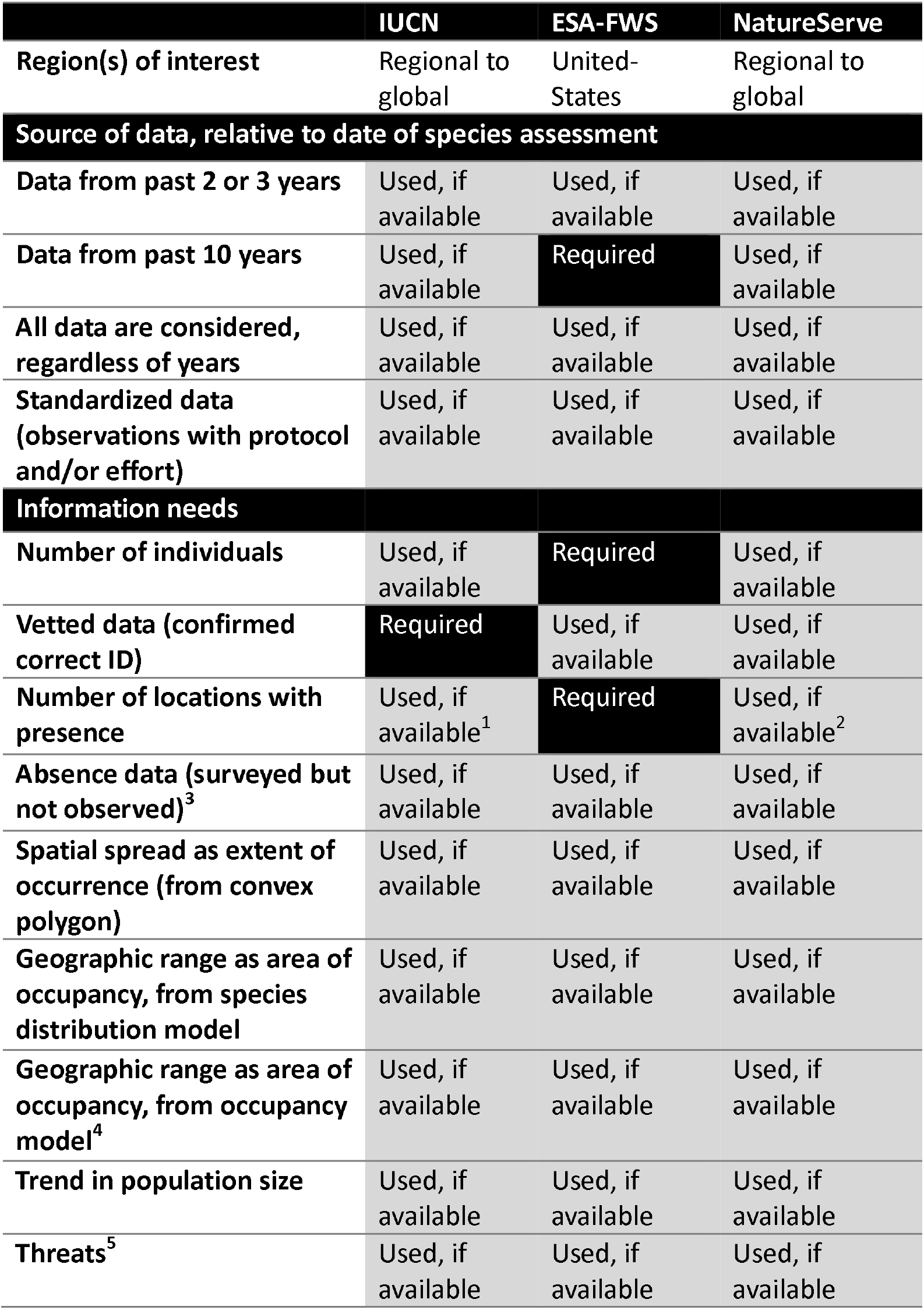
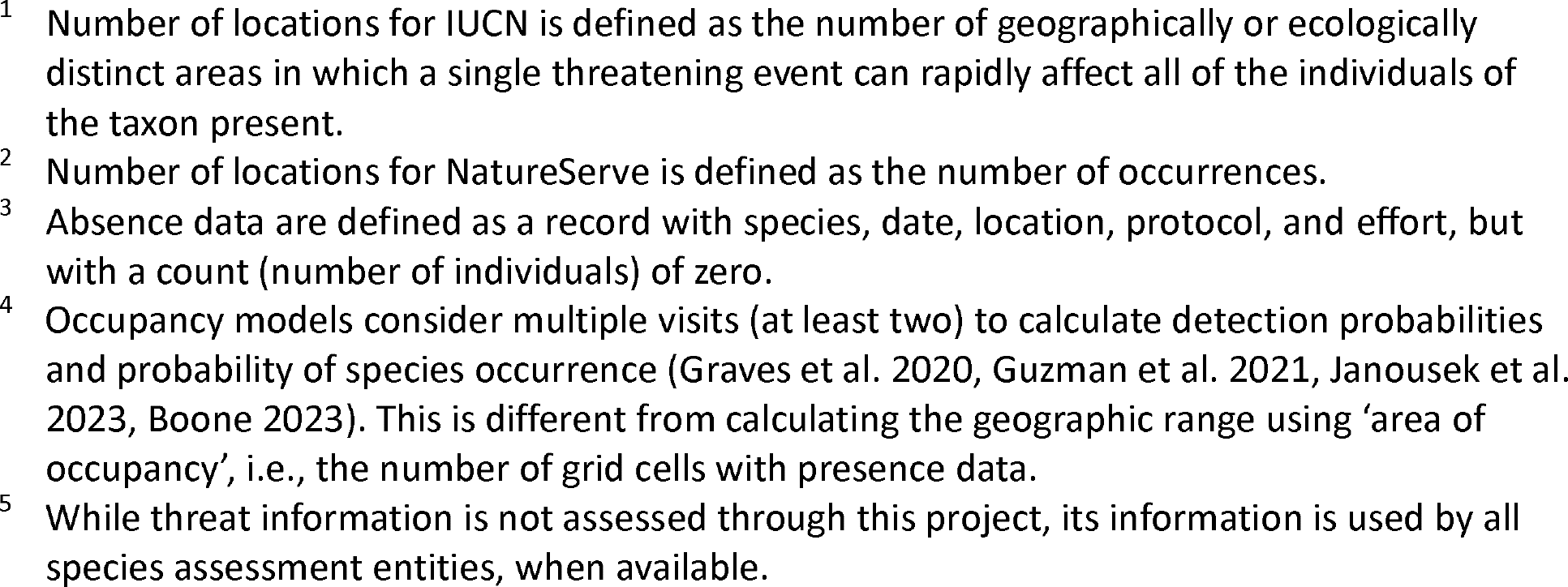
List of major entities/unions completing species assessments for conservation status for species located in the United-States – International Union for Conservation of Nature (IUCN), Endangered Species Act by U.S. Fish & Wildlife Service (ESA-FWS), and the NatureServe Network – and the source of data and information needs associated with each.

|  | **IUCN** | **ESA-FWS** | **NatureServe** |
| --- | --- | --- | --- |
| **Region(s) of interest** | Regional to global | United-States | Regional to global |
| **Source of data, relative to date of species assessment** |  |  |  |
| **Data from past 2 or 3 years** | Used, if available | Used, if available | Used, if available |
| **Data from past 10 years** | Used, if available | **Required** | Used, if available |
| **All data are considered, regardless of years** | Used, if available | Used, if available | Used, if available |
| **Standardized data (observations with protocol and/or effort)** | Used, if available | Used, if available | Used, if available |
| **Information needs** |  |  |  |
| **Number of individuals** | Used, if available | **Required** | Used, if available |
| **Vetted data (confirmed correct ID)** | **Required** | Used, if available | Used, if available |
| **Number of locations with presence** | Used, if available<sup>1</sup> | **Required** | Used, if available<sup>2</sup> |
| **Absence data (surveyed but not observed)<sup>3</sup>** | Used, if available | Used, if available | Used, if available |
| **Spatial spread as extent of occurrence (from convex polygon)** | Used, if available | Used, if available | Used, if available |
| **Geographic range as area of occupancy, from species distribution model** | Used, if available | Used, if available | Used, if available |
| **Geographic range as area of occupancy, from occupancy model<sup>4</sup>** | Used, if available | Used, if available | Used, if available |
| **Trend in population size** | Used, if available | Used, if available | Used, if available |
| **Threats<sup>5</sup>** | Used, if available | Used, if available | Used, if available |

- <sup>1</sup> Number of locations for IUCN is defined as the number of geographically or ecologically distinct areas in which a single threatening event can rapidly affect all of the individuals of the taxon present. - <sup>2</sup> Number of locations for NatureServe is defined as the number of occurrences. - <sup>3</sup> Absence data are defined as a record with species, date, location, protocol, and effort, but with a count (number of individuals) of zero. - <sup>4</sup> Occupancy models consider multiple visits (at least two) to calculate detection probabilities and probability of species occurrence (Graves et al. 2020, Guzman et al. 2021, Janousek et al. 2023, Boone 2023). This is different from calculating the geographic range using 'area of occupancy', i.e., the number of grid cells with presence data. - <sup>5</sup> While threat information is not assessed through this project, its information is used by all species assessment entities, when available.

We used publicly-available biodiversity data for wild bees to evaluate the quality and quantity of U.S. bee data, specifically in the context of understanding data availability for higher-quality status assessments. With an eye towards reproducibility and the needs of conservationists who perform species assessments, we focused on the propensity to report much-needed metadata such as sampling methods (protocol and effort) for data collection. Several recent studies have examined the status of bee data at broad spatial (U.S. and global) and taxonomic scales and used these data to detect trends in patterns such as data gaps, species richness, and species ranges (Orr et al., 2021; Zattara and Aizen, 2021; Chesshire et al., 2023). Our work complements these studies by extending the focus to the quality of these data with respect to reporting, reproducible data collection, and the interests of conservation decision-makers. Based on the recognition that there are different levels and types of data quality, that data are often lacking for bees compared to many other taxa, and that species assessments are valuable even when performed with minimal available data, we used a flexible approach where we examined how many bee species would have sufficient quantity and quality of data to meet different analysis quality thresholds. We found that even with relatively relaxed criteria, there are major data and reporting gaps that hinder bee species status assessments, and thus are ultimately limiting bee conservation action in the U.S. In particular, information about sampling protocol and effort is very rarely reported in publicly available data sets, which limits the ability to replicate data collection methods and use data for more rigorous analyses to assess status and trends. In spite of these limitations, we detected a general pattern where more than half of U.S. bee species have not been observed in the last decade of our dataset (i.e. from 2012-2021). These species are important candidates for future targeted data collection efforts and can serve as an initial list of bee species that may be of conservation concern for consideration by states in conservation planning (supplementary material S1; Figure 3). Moving forward, we provide suggestions for improving data reporting for bee conservation and provide resources to aid states and other conservation practitioners in their efforts to conserve bee species.

## MATERIALS AND METHODS

We evaluated the extent to which the current state of bee data is sufficient to support rigorous assessments of species for conservation status based on population size, distribution, and trend analyses (Table 1). Species conservation status assessments (IUCN, 2012; Master et al., 2012; U.S. Fish and Wildlife Service, 2016) are instrumental in prioritizing species for conservation action, and the data generated from them contribute to management and recovery plans (e.g., U.S. Fish and Wildlife Service, 2021). As such, we focus on data requirements to improve species conservation status assessments. Some of these requirements are not readily available and were identified through a series of communications with the various entities completing these assessments (see acknowledgment section; Table 1).

### Creation of dataset

We began our analyses with the bee dataset compiled by Chesshire et al. (2023), which was downloaded from the Global Biodiversity Information Facility (GBIF; https://www.gbif.org/) and Symbiota Collections of Arthropods Network (SCAN; https://scan-bugs.org/) in February 2021. This dataset contains 1.9 million records observed from 1700 through early 2021. We then supplemented the Chesshire et al. (2023) dataset with 111,216 records observed in 2021 and downloaded from GBIF (GBIF.org, 2022) and SCAN (SCAN, 2022) in August 2022. We applied a series of filters to the 2021 dataset following the process performed by Chesshire et al. (2023). First, we confirmed the 2021 species validity through expert assessment; this included updating species names containing typographical errors or names that were taxonomically revised, and removing species lacking species-level identification or not reliably confirmed to be present in the U.S. Next, we removed all records of honey bees (*Apis mellifera*) from the 2021 data, as this is a managed or feral species in the U.S, rather than a wild native species. Last, we removed records that were outside the contiguous U.S. and those for which the uncertainty about a location exceeded 15 km. We recognize that some records with a high uncertainty may be rare species whose locations were obscured to protect their location. After filtration, the 2021 data contributed an additional 68,026 records, resulting in a total of 1,991,840 records. We would like to acknowledge the following institutions, whose data contributed to at least five percent of all records used in the analysis: American Museum of Natural History (Johnson, 2020), iNaturalist (iNaturalist contributors and iNaturalist, 2023), University of Kansas Biodiversity Institute (Bentley and Thomas, 2023), U.S. Department of Agriculture (Ikerd, 2019), U.S. Geological Survey (Droege and Maffei, 2023). All analyses were completed in R (R Core Team, 2023).

### Data suitability for conservation status assessments

To evaluate the suitability of records in our dataset for conservation status assessments, we established two classes of records: (1) Complete records, which included detailed information about species nomenclature, date (month, day, year), location (latitude, longitude, location uncertainty), count (number of bees per record), sampling protocol (e.g., trap type or protocol name), and sampling effort (e.g., number of traps/observers and sampling duration), and can be used to conduct population sizes and trend estimates, and (2) Partially complete records, which identified species and location within 15 km certainty but were missing other information. A record was determined to provide sampling protocol if it included any information about how the associated bee was collected. We categorized any provided sampling protocol information into the following groups: net (hand-netted), pan (pan trap), net and pan, malaise (malaise trap), or other traps (which included, for example, vane, pitfall, and light traps). Similarly, a record was determined to provide sampling effort if it had any information related to number of traps, number of collectors, and/or sampling duration.

### Species summaries and data thresholds

To investigate the spatial patterns of bee occurrences across the contiguous U.S., we summarized the number of bee records within 25 km apothem (inradius) hexagon grid cells covering this region. We created species summaries by calculating the total number of records per species and the number of decades during which they were sampled. We also assessed the number of bee records per species from 2012 to 2021, the associated number of records per trap type, number of unique locations, and the number of records with sampling protocol and/or effort information. (supplementary material S2). The most recent ten-year window was selected because it represents a period of interest for several entities completing species assessments (IUCN, 2012; Master et al., 2012) and aligns with the ten-year cycle of updating Species of Greatest Conservation Needs lists for State Wildlife Action Plans (Mawdsley and Humpert, 2016; deMaynadier et al., 2023).

Because species assessments typically include details about geographical spread and range, such as the extent of a species occurrence and area of occupancy, we established whether each species met a set of progressively more stringent hierarchical data thresholds. Each additional threshold allows for a more detailed assessment of the species distribution, most of which are performed, if possible, by the major entities that perform bee species status assessments (Table 1). The thresholds presented here are minimum requirements that are dramatically reduced relative to the norms used for vertebrate species (Mackenzie and Royle, 2005; Devarajan et al., 2020; Johnston et al., 2021); we have modified them to account for some of the additional challenges of collecting bee data and focus on the relative number of species for which it would be possible to complete each level of analysis. We tested the sufficiency of each species’ data for the following thresholds using the 2012-2021 dataset:

1. Convex polygon requirements: three unique locations, which is the minimum requirement for this calculation.
2. Minimum species distribution model requirements: 30 records and 30 unique locations (Stockwell and Peterson, 2002; Wisz et al., 2008; Luan et al., 2020).
3. Low-resolution occupancy model requirements: Using a 100 km hexagonal cells grid (Jackson et al., 2022), we selected all cells with at least two visits, where a visit was defined by a unique combination of date and location. All species present in at least 30 cells satisfied this threshold.
4. Higher resolution occupancy model requirements: Using a 10 km hexagonal cells grid (Janousek et al., 2023) and records containing protocol (trap type) information, and selecting all cells with at least two visits. All species present in at least 30 cells satisfied this threshold.

Occupancy models, when completed using best practices, provide a more accurate assessment of a species distribution because they consider imperfect detection (Graves et al., 2020; Guzman et al., 2021; Boone et al., 2023b, 2023a; Janousek et al., 2023; Otto et al., 2023). Their creation requires information about surveys where species were detected or not detected (i.e., absence data where the individual count is zero), and multiple visits at the same site.

### State-specific analyses

To assess the number of bee species for each state in the contiguous U.S. over time, we summarized our dataset over all years (1700-2021) and in a recent decade (2012-2021). We assessed the number of bee species ever observed in each state and determined the percentage of these that were recently observed (2012 to 2021). We also identified the species in each state that had been previously recorded but were not observed from 2012 to 2021 (supplementary material S1).

## RESULTS

### Availability of US bee data

Data downloaded from public repositories including GBIF and SCAN often require extensive data cleaning for analytical purposes, substantially reducing the amount of data available for conservation-related analyses, such as species assessments (Chesshire et al. 2023, this paper). Nearly 25% of the records in the Chesshire et al. (2023) dataset were discarded because they lacked species or location (personal communication, Paige Chesshire). Bee record quality and quantity have improved over the years, particularly since the 2000s (Figure 1). Yet, only a small fraction of publicly available records were complete and contained information about protocol and effort (n = 733, or 0.04% of all records; supplementary material S2); all of these records were from 2021. Most records (92%) collected from 2019 to 2021 were submitted through iNaturalist and lacked sampling protocol and effort information. Complete records provide the data required to estimate population size and trend, as they enable comparisons of bee abundance and richness over time and space.

**Figure 1.**
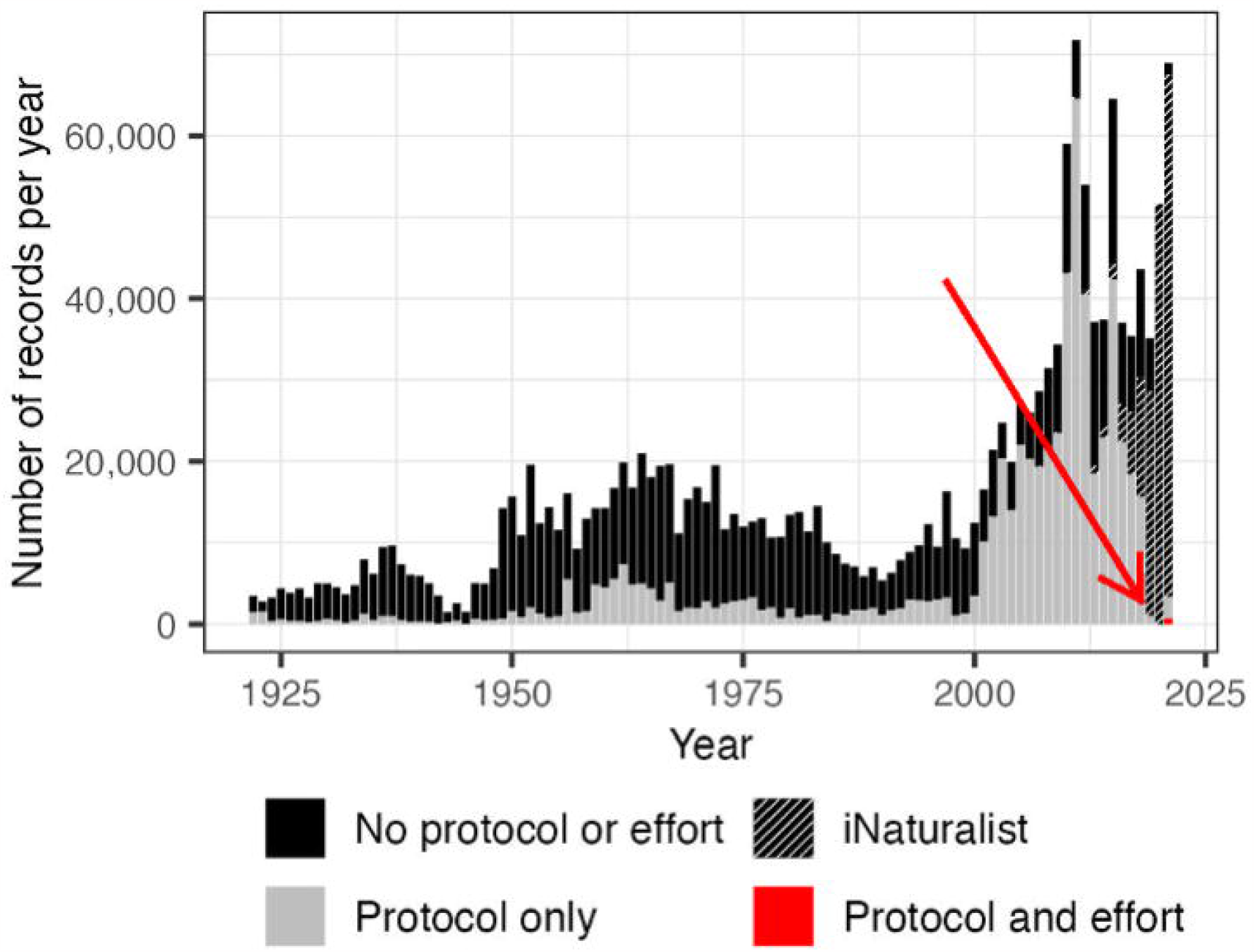
Number of bee records per year, for 1922 to 2021 (few records exist prior to 1922; n = 73,695), where the black bar section represents the number of records without protocol (trap type) nor effort (duration and/or number of traps/volunteers), gray represents records with only protocol information, and red represents *complete* records containing both protocol and effort information.

With respect to data availability per species, particularly for species assessments, only 33% of the 3,219 species recorded in the contiguous U.S. had sufficient data to describe their geographic spread using convex polygons (Table 2; supplementary material S2). Few species had sufficient data to generate distribution models (11%), lower-quality occupancy models (6%), or higher-quality occupancy models (5%).

**Table 2.** The number and associated percentage of bee species for which we have records from 2012-2021 and meet the data thresholds for four geographical range analyses. Total number of species considered is 3,219. “Threshold” refers to the minimum data needs for the analysis listed.

| **Time period or analysis** | **Threshold** | **Number of species** | **Percent of species** |
| --- | --- | --- | --- |
| **2012 to 2021** | Number of species for which we have at least one record. | 1,528 | 47.4% |
| **Convex polygon – Records from 2012 to 2021** | Number of species with at least 3 unique locations | 1,070 | 33.2% |
| **Species distribution model – Records from 2012 to 2021** | Number of species with at least 30 records and 30 unique locations | 369 | 11.5% |
| **Lower-quality occupancy model – Records from 2012 to 2021** | Selected 100 km grid cells with at least two surveys. Number of species present in at least 30 grid cells. | 194 | 6.0% |
| **Higher-quality occupancy model – Records from 2012 to 2021** | Selected 10 km grid cells with at least two surveys, using only records with trap type information. Number of species present in at least 30 grid cells. | 148 | 4.6% |

### Spatial distribution of bee data

Summarizing our dataset within 25 km hexagon grid cells revealed a greater availability of data on the west and east coasts, with sparse coverage in many parts of the interior U.S. (Figure 2). The percentage of grid cells without bee records, represented by the white surface in Figure 2, ranged from 6% across all years combined, to 23% from 2012 to 2021. The majority of grid cells contained fewer than 100 records, whether across all years (57%) or from 2012-2021 (86%).

**Figure 2.**
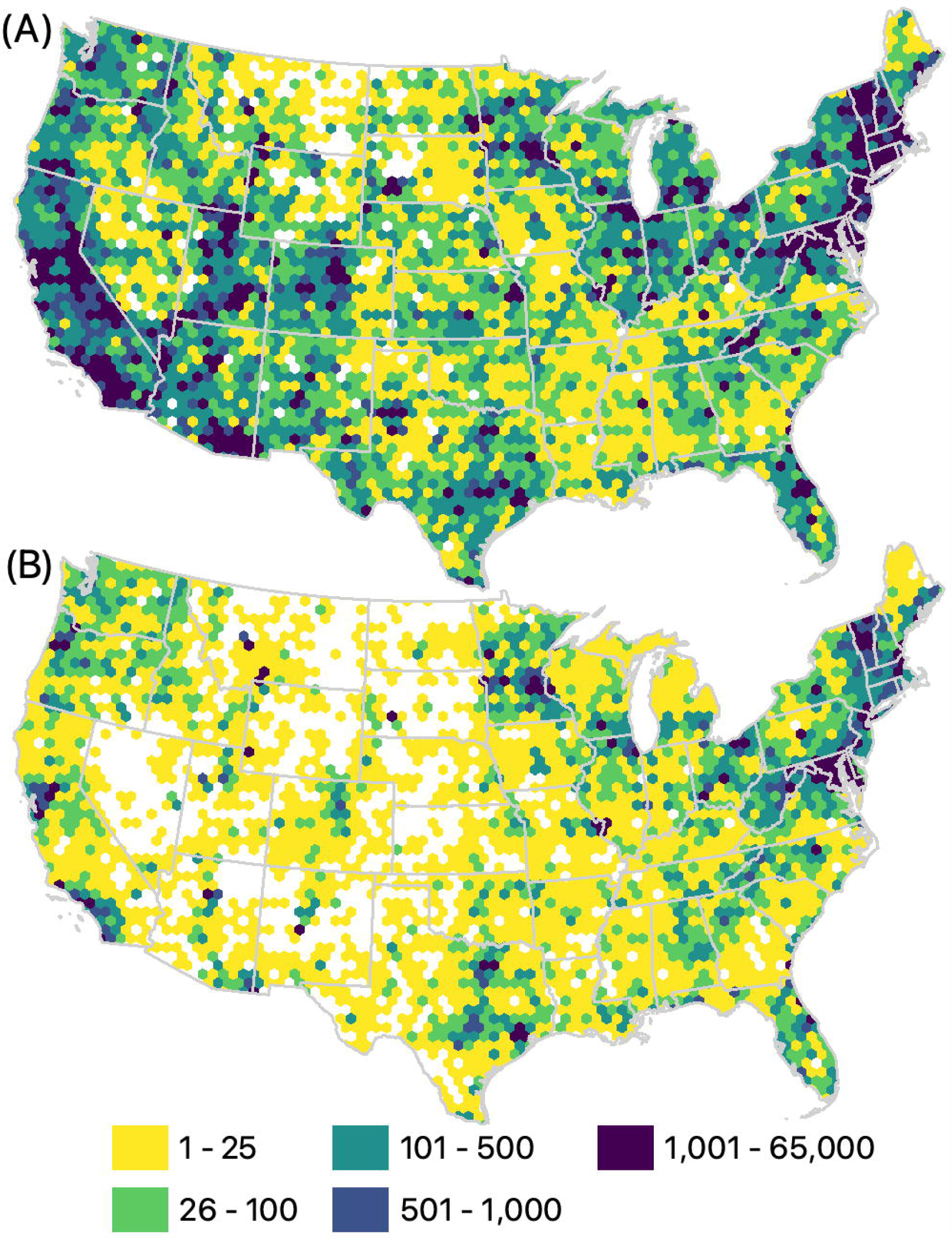
Number of bee records per 25 km hexagon grid cell. (A) Total number of records available since 1700 (n = 1,991,840). (B) Number of records from 2012 to 2021 (n = 464,845).

**Figure 3.**
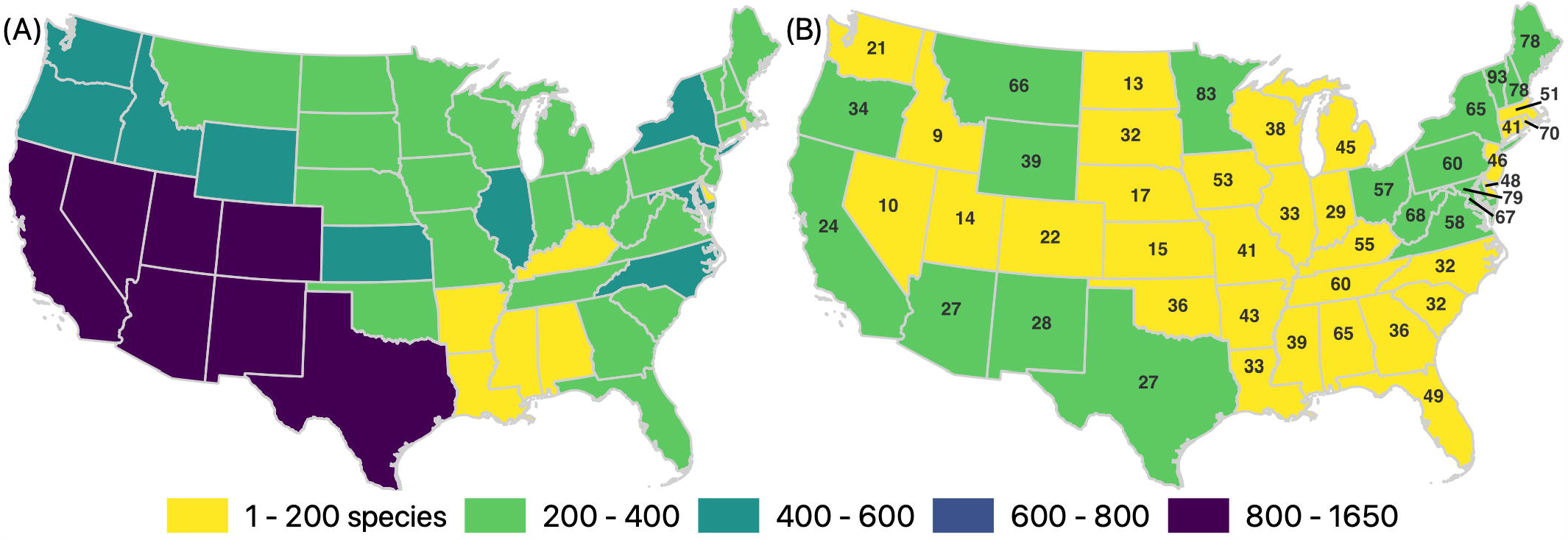
Number of species sampled per state and time period. (A) Total number of species observed across all years (1700-2021), per state (B) The colors represent the number of species observed from 2012 to 2021, and the numbers overlaid on each state represent the percentage of the total number of species known from each state that were observed during the recent time period. The data associated with these maps, including a list of the species that have been observed in each state but were not observed from 2012-2021, are available in the supplementary material SI.

Fewer than half (47%) of the 3,219 bee species were recorded from 2012 to 2021 (Table 2; supplementary material S2). Only 5 states had >75% of their known U.S. bee species recorded between 2012 and 2021, whereas 30 states had fewer than half (supplementary material S1; Figure 3). Most species (3,003 species) that had been observed prior to 2012 were no longer observed between 2012-2021 in at least one state in which they historically occurred.

## DISCUSSION

As populations of wild bees continue to decline, a broad community of scientists, practitioners, and members of the public have begun to galvanize around the need to better track species status and trends and take action to conserve bee populations (National Research Council, 2007; Pollinator Health Task Force, 2015; Mawdsley and Humpert, 2016; Inouye et al., 2017; Woodard et al., 2020; deMaynadier et al., 2023). Because the availability of high-quality data remains a limiting factor to conservation, we evaluated the extent to which publicly available data are sufficient to support species-level assessments of population status and trends. Based on the most minimal data standards (e.g., specifying location, sampling protocol, effort), we found that only a small fraction of data records are suitable for use in species assessments.

Our analysis points to a striking paucity of complete records in public data repositories for wild bees, and this greatly limits our ability to assess population status and trend for most species.

Many records were excluded because species and/or location information was not provided or was not precise enough, often because the records originated from older collections or, in terms of location uncertainty, because data providers chose to not make this information publicly available. Most recent data (92% of records from 2019 through 2021) come from iNaturalist, which is a publicly available platform to which volunteers can submit photograph-based observations, often opportunistically-obtained. iNaturalist data are heavily biased with respect to their geographic distribution and photographed species, with more records from areas of higher human habitation or use and representing larger bee species. iNaturalist records were also seldom complete because they failed to include information on sampling effort or protocol, which may reflect how the screens to enter sampling information are not obvious to an observer. Despite improving submission rates and record completeness over the last two decades, our ability to collect complete data on wild bees continues to fall short of what is sorely needed by the scientific and conservation communities. Case in point, information about sampling protocol and effort was reported for only 0.04% of records in the dataset. All complete records are from 2021, the last year represented in our dataset. Based on our species-specific examination, we also found a striking lack of data that met the core criteria for species assessments, outlined in Table 1.

Insufficient data existed to describe geographic spread (67% of species), generate distribution models (89%), and support either lower- (94%) or higher-quality (95%) occupancy models.

Our focus was specifically on understanding what proportion of publicly available bee data are complete with respect to reporting information, such as protocol and effort, that is required for many analyses of species status and trends. We note, however, that there are special considerations for bee data collection and curation that influence the quantity and quality of available data. The bee data collection community faces obstacles such as the complexities of sampling a diverse group of small and highly mobile species, taxonomic challenges that are exacerbated by lack of funding and support (Gonzalez et al., 2013; Woodard et al., 2020), and bee identification and digitization backlogs. These challenges are being met (Cobb et al., 2019; Seltmann et al., 2021; Chesshire et al., 2023; Dorey et al., 2023) but have almost certainly contributed to a relative lack of data to date. We also note that some major efforts to collect bee data across the U.S., such as through the USGS Native Bee Inventory and Monitoring Program, have only recently uploaded a complete version of their records, while others, such as state atlas projects, will yield more, and higher-quality, data in the coming years.

One worrisome finding was that most bee species in the U.S. in our data set were recorded only prior to 2012. Despite an exponential increase in bee data collection since 2012, more than half of bee species have not been recorded, at least based on data housed within major public repositories. One explanation for this pattern is that local extinctions of species have gone unnoticed in well-surveyed areas. More broadly, this would be consistent with the idea that more bee species are declining than are currently recognized (Zattara and Aizen, 2021), and certainly more than are currently being protected at the state and Federal levels. An alternative explanation is that these species have been observed since 2012, but these records are not housed in major public biodiversity data repositories. They may exist instead in privatized data collections; in some cases, these collections may even be especially likely to contain data for at-risk species.

These species may have also been observed in recent surveys, but these data are simply not yet publicly available. This highlights the need, when possible in light of restrictions in data sharing, for statewide wild bee atlases and other bee data collection efforts to fully share their data on public repositories. This facilitates data sharing, openness, and reproducibility, and allows for species assessments and other analyses to be performed by the broader bee research and conservation communities, to the overall benefit of bee conservation. Regardless of which of the two aforementioned scenarios is true, we recommend that species that have not been observed in our data set in the most recent decade (2012-2021; see lists by state in supplementary materials S1 and S2) be candidates for more targeted data mining and collection. This list of putatively “missing” species, if they are confirmed absent in all available data sets, may also be used to inform statewide conservation planning. This could include their placement on lists of Species of Greatest Conservation Need (Mawdsley and Humpert, 2016; deMaynadier et al., 2023), as the bee species that have not been recently observed may indeed be of great conservation concern.

Moving forward, we see several actions that could improve the quality of wild bee data, many of which center on greater standardization of data collection, reporting, and management. Data standards, if they have been applied at all, have been applied differently across the myriad bee survey efforts that have taken place over the past two decades (Montgomery et al., 2020, 2021; Woodard et al., 2020). Concrete data standards and best practices would help ensure that we have data we need to complete robust, accurate assessments. We suggest encouraging data collection that adheres to an existing standard (e.g., Darwin Core) and promotes FAIR (findable, accessible, interoperable, reusable) data sharing principles (Wieczorek et al., 2012; Wilkinson et al., 2016). We further suggest defining a standardized vocabulary of accepted entries for each data variable collected, as was proposed by Montgomery et al. (2021). This will facilitate data sharing, interoperability, and use, as the terms and the corresponding entries will align across multiple datasets. We provide a list of suggested terms in the supplementary material (S3) that describe the date, location, count, species nomenclature, sampling protocol, and sampling effort associated with the data. The significant gaps we, and others (Chesshire et al., 2023), have identified in wild bee data and the potential for wild bee species declines or losses compel us to request that wild bee data collectors consider integrating our suggested changes in data collection and reporting into their current and future inventory, survey, and monitoring efforts. Doing so would contribute to the creation of quantitatively supported, sound wild bee conservation policy and practice, which is critical in the protection of these species. Combined, the protocol and effort fields allow us to compare bee abundance and richness across time and locations, which is essential for estimating accurate trends. Data collected now using these methodologies would enable calculation of trends in just a few years.

## CONCLUSIONS

Despite the accumulation of decades worth of wild bee data in public repositories, we found that the quality of available records is often insufficient to support the rigorous estimation of range/distribution, population size, trends, or other information needed for species assessments. This is a pattern that is generally observed for invertebrates, not only bees. The shortcomings that we outline in this paper can be readily addressed with improvements in data collection and reporting, along with the use of more standardized protocols. With coordinated outreach and education to improve data quality, we can build capacity in the broad network of scientists and practitioners working to identify species most in need of conservation, elucidate potential drivers of decline, and guide strategic action to halt wild bee declines.

## FUNDING

The research included in this paper was made possible through funding by the Walmart Foundation to JSR, ADR, and AJ. The findings, conclusions and recommendations presented in it are those of the authors alone, and do not necessarily reflect the opinions of the Walmart Foundation. The USDA National Institute of Food and Agriculture provided funding for SHW (USDA NIFA 2020-67014-31865).

## ACKNOWLEDGMENTS

Thanks to the following for providing information about species assessments on behalf of their entities (names are in alphabetical order of last name): Jeff Everett (US Fish & Wildlife Service), Rich Hatfield (Xerces Society), Eric Miskow (Nevada Division of Natural Heritage), Kristin Szabo (Nevada Division of Natural Heritage), Matthew Schlesinger (New York Natural Heritage Program), Anna Walker (New Mexico BioPark Society), Erin White (New York Natural Heritage Program), Paul Williams (Natural History Museum), and Bruce Young (NatureServe). Thanks to Paige Chesshire for sharing information about the quality control and filters that were applied to the Chesshire et al. (2023) dataset. Thanks to Melissa Guzman for sharing how occupancy models can (and cannot yet) be used to answer questions about the status and trends of bee populations. We appreciate the feedback from Leif Richardson (Xerces Society for Invertebrate Conservation) and Mark Buckner (Cornell University), which helped improve our paper. Thanks to the following data owners, which provided at least 5% of the records used in the dataset: American Museum of Natural History, iNaturalist, University of Kansas Biodiversity Institute, U.S. Department of Agriculture, and U.S. Geological Survey; we also cite the providers associated with these collections in our references. Lastly, but certainly not least, we want to express our gratitude to the Cornell Atkinson Center for Sustainability, particularly Patrick Beary and Gail Phillips, for their collaboration with ADR, JSR, and AJ, throughout the project and their valuable input in securing funding and maintaining a positive relationship with our funder.

## Notes

### Competing Interest Statement

The authors have declared no competing interest.

